# Evolution of vertebrate gill covers via shifts in an ancient Pou3f3 enhancer

**DOI:** 10.1101/2020.01.27.918193

**Authors:** Lindsey Barske, Peter Fabian, Christine Hirschberger, David Jandzik, Tyler Square, Pengfei Xu, Nellie Nelson, Haoze Vincent Yu, Daniel M. Medeiros, J. Andrew Gillis, J. Gage Crump

## Abstract

Whereas the gill chambers of extant jawless vertebrates (lampreys and hagfish) open directly into the environment, jawed vertebrates evolved skeletal appendages that promote the unidirectional flow of oxygenated water over the gills. A major anatomical difference between the two jawed vertebrate lineages is the presence of a single large gill cover in bony fishes versus separate covers for each gill chamber in cartilaginous fishes. Here we find that these divergent gill cover patterns correlate with the pharyngeal arch expression of Pou3f3 orthologs. We identify a Pou3f3 arch enhancer that is deeply conserved from cartilaginous fish through humans but undetectable in lampreys, with minor sequence differences in the bony versus cartilaginous fish enhancers driving the corresponding single versus multiple gill arch expression patterns. In zebrafish, loss of Pou3f3 gene function disrupts gill cover formation, and forced expression of Pou3f3b in the gill arches generates ectopic skeletal elements resembling the multiple gill cover pattern of cartilaginous fishes. Emergence of this Pou3f3 enhancer >430 mya and subsequent modifications may thus have contributed to the acquisition and diversification of gill covers and respiratory strategies during gnathostome evolution.

## Main text

Jawed vertebrates have evolved different anatomical structures to protect and ventilate their gills (Fig. 1a,g). In bony fishes, the second pharyngeal arch (hyoid) grows caudally over the posterior gill-bearing arches to form a single large gill cover (operculum) supported by intramembranous bones. Although humans and other tetrapods lack functional gills, growth of the hyoid arch over the posterior arches is conserved. The hyoid outgrowth ultimately merges with the trunk to close off the pharyngeal cavity, with defects in this process resulting in branchial cysts and fistulae in humans (e.g. Branchiootorenal syndrome)^1, 2^. By contrast, in elasmobranch cartilaginous fishes (sharks, skates, and rays), the hyoid and four posterior arches each undergo posterior-directed growth to generate five separate gill covers supported by cartilaginous rays. In holocephalans, the other extant group of cartilaginous fishes, five separate gill covers also form initially, yet subsequent stalled growth of the four posterior covers results in a single hyoid-derived gill cover^3^. Gill cover formation is stimulated by an epithelial signaling center called the posterior ectodermal margin (PEM) that develops along the caudal rim of the hyoid arch in bony fishes and arches 2-6 in cartilaginous fishes (later refined to just arch 2 in holocephalans) (Fig. 1g)^1, 3, 4^. The PEM is a source of Shh, Fgfs, and Bmps^4–7^ that are believed to act on the underlying neural crest-derived mesenchyme that generates the gill cover skeleton and connective tissues. Whether this mesenchyme possesses a pre-pattern for single versus multiple gill covers, or simply receives instructive information from the PEM, had not been examined.

**Fig. 1.**
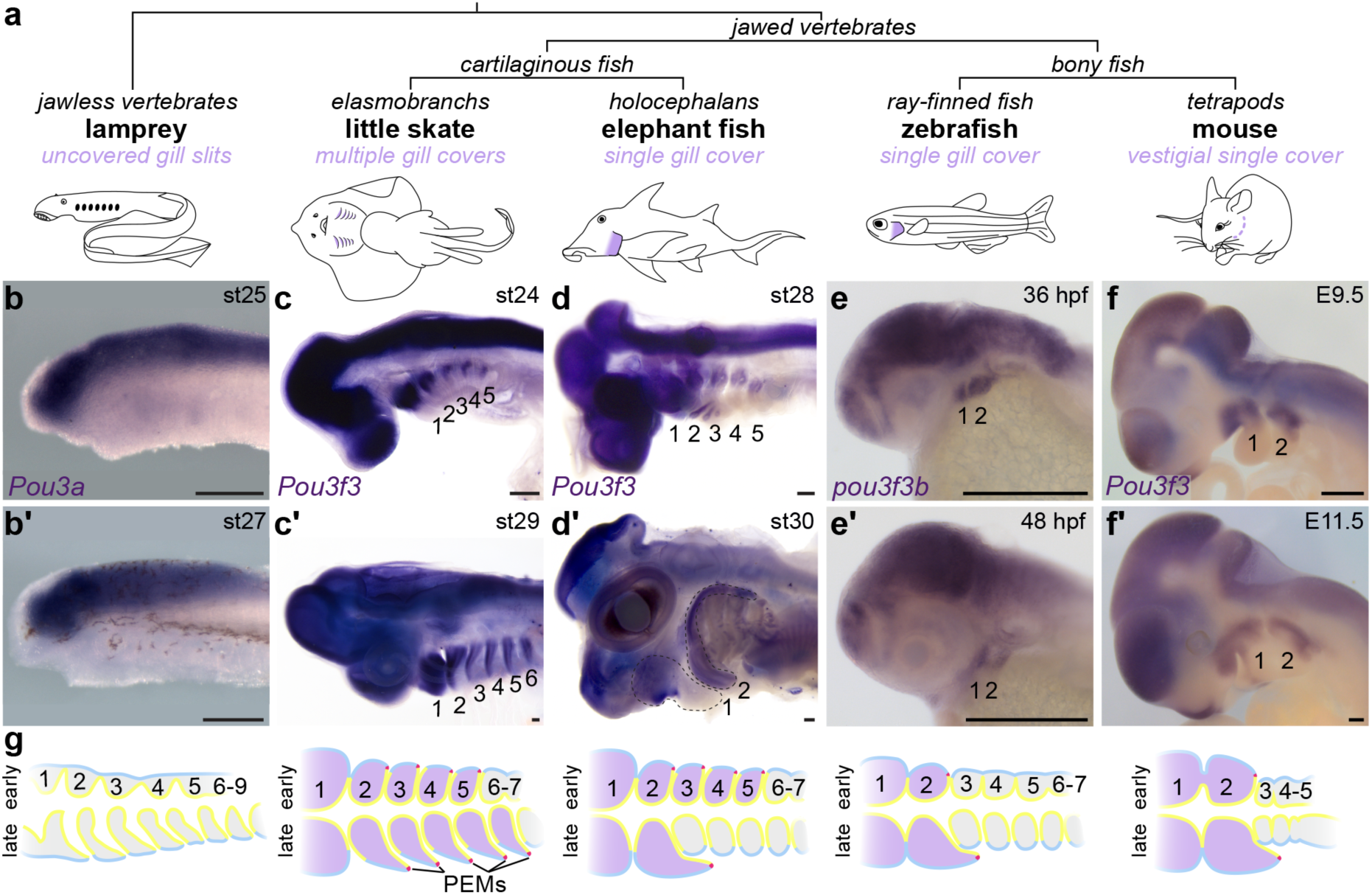
Arch Pou3f3 expression correlates with gill cover formation. **a**, Vertebrate phylogeny highlighting the gill cover type of each lineage. **b-f**, In situ hybridizations for Pou3f3 homologs in lamprey, little skate, elephant fish, zebrafish, and mouse show conserved expression in the CNS but divergent patterns in the pharyngeal arches. Note the later suppression of Pou3f3 in the posterior arches of the elephant fish. The dotted lines in **d**’ outline the first and second arches. Scale bars = 250 μm. **g,** Schematized frontal sections through the pharyngeal arches (numbered) at early (top) and late (bottom) stages, with arch Pou3f3 expression coded in purple, PEMs in red, neural crest-derived mesenchyme in grey, endoderm in yellow, and ectoderm in blue.

POU Class 3 Homeobox 3 (POU3F3), also known as BRN1, is a proneural transcription factor with widespread expression in the developing central nervous system^8^ as well as in dorsal pharyngeal arch mesenchyme. We noted that zebrafish^9^, mouse^10^, and frog^11^ Pou3f3 homologs are atypical among arch-enriched genes in that they are expressed only in the mandibular (first) and hyoid arches and not in the more posterior gill-forming arches (Fig. 1e-f; compare with *dlx2a*, *dlx6a*, and *hand2* in Extended Data Fig. 1a-b). At later stages, zebrafish *pou3f3a* and *pou3f3b* are restricted to the growing gill cover (Extended Data Fig. 1c-d). This restriction to the forming gill cover prompted us to examine the expression of *Pou3f3* orthologs in species with divergent gill cover patterns. In the sea lamprey (*Petromyzon marinus*), a jawless vertebrate lacking gill covers, we detected expression of the three Pou3 homologs *Pou3a*, *Pou3b*, and *Pou3c* in the brain but not the pharyngeal arches (Fig. 1b, Extended Data Fig. 2). In the little skate (*Leucoraja erinacea*), an elasmobranch with five gill covers, *Pou3f3* is expressed in the brain and arches 1-6 (Fig. 1c). In the elephant fish (*Callorhinchus milii*), a holocephalan, *Pou3f3* initially displayed expression in arches 1-5 and the brain but later became restricted to the hyoid and mandibular arches, consistent with the secondary regression of the posterior covers^3^ (Fig. 1d). These patterns collectively indicate ancestral expression of *Pou3f3* homologs in the brain, with co-option of *Pou3f3* into arch mesenchyme in jawed vertebrates tightly correlating with the presence, number, and developmental progression of gill covers across diverse species.

We next sought to identify cis-regulatory elements that could explain the distinct Pou3f3 expression patterns. Assay for transposase-accessible chromatin sequencing (ATAC-seq) on sorted zebrafish *sox10*:DsRed^+^; *fli1a*:EGFP^+^ arch mesenchyme versus control double-negative cells revealed differentially accessible, non-orthologous regions downstream of *pou3f3a* (Dr4, 380 bp) and *pou3f3b* (Dr1, 2033 bp) (Fig. 2a, Extended Data Fig. 3a; Extended Data Table 1). In transgenesis assays, both *pou3f3b*-Dr1 and *pou3f3a*-Dr4 drove GFP expression in neural crest-derived mesenchyme of the dorsal mandibular and hyoid arches at 36 hours post fertilization (hpf), with limited or no expression in the posterior arches (Fig. 2c-d; Extended Data Fig. 3c). Neither Dr1 nor Dr4 drove CNS expression, in contrast to a *pou3f3b^Gal4ff^*; *UAS:*nlsGFP knock-in reporter line that recapitulates endogenous *pou3f3b* expression in the CNS as well as the dorsal first and second arches (Fig. 2d, Extended Data Fig. 4a-c). At 6-7 days post fertilization (dpf), *pou3f3b*-Dr1 but not *pou3f3a*-Dr4 continued to drive robust expression in the operculum (Fig. 2e, Extended Data Fig. 3c), consistent with the broader endogenous expression of *pou3f3b* during opercular outgrowth (Extended Data Fig. 1c-d). We focused on *pou3f3b*-Dr1 for further analysis, as *pou3f3a*-Dr4 is conserved only in the teleost fish species most closely related to zebrafish (e.g. cavefish, catfish, and piranha) (Extended Data Fig. 5). Within *pou3f3b*-Dr1, we identified a 515-bp 5’ fragment (“Dr1A”) that drove expression in only the maxillary domain, similar to *pou3f3a*-Dr4 (Extended Data Fig. 3b-d). We also identified a central 610-bp sequence (“Dr1B”) that drove robust expression in the dorsal mandibular and hyoid arches starting at 36 hpf and the growing operculum at 7 dpf (Fig. 2a, d-e). Whereas Dr1A is found only in teleost fish (n = 43/45 genomes; Extended Data Fig. 5), Dr1B is exceptionally well conserved across gnathostomes (Fig. 2b). Homologous sequences were recovered in syntenic regions of all mammalian (n = 20), avian (5), reptilian (5), amphibian (1) and cartilaginous fish (4) genomes evaluated (Extended Data Table 1). Zebrafish Dr1B was the most divergent in an alignment of 11 vertebrate 1B sequences, and no 1B sequence could be retrieved in 12/45 teleosts (Extended Data Figs. 5-6), in line with previous work showing rapid evolution of regulatory sequences following the teleost whole-genome duplication ∼300 mya^12^. We also did not identify homologous Dr1B sequences in lamprey or two invertebrate chordates (amphioxus and tunicate) that lack gill covers. In addition, we independently predicted the human “Hs1B” and mouse “Mm1B” sequences as enhancers by virtue of their open chromatin and 25-state epigenomic annotation in published datasets for human and mouse neural crest-derived cells (Fig. 2a)^13–16^.

**Fig. 2.**
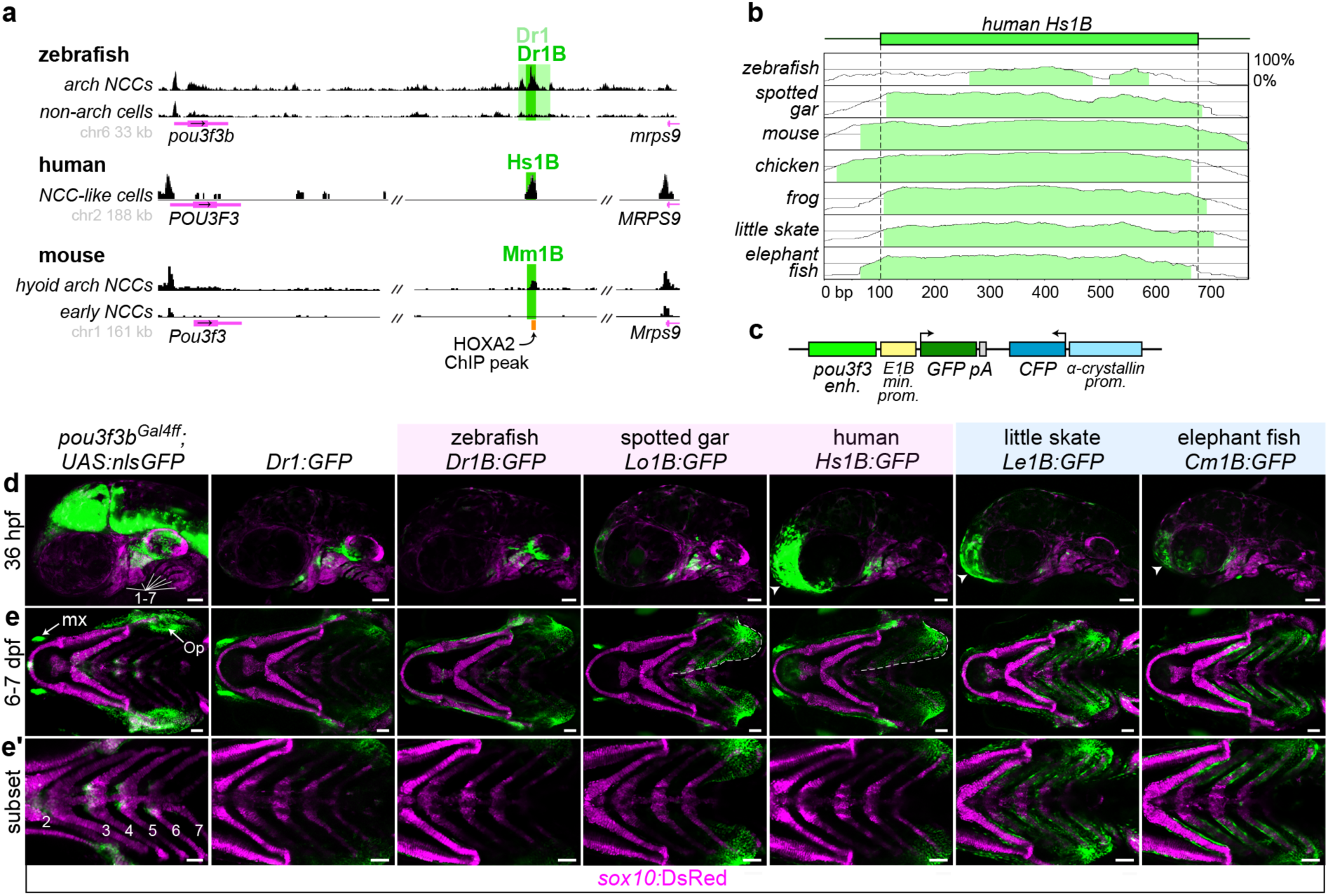
A deeply conserved Pou3f3 enhancer differentially regulates arch expression in bony and cartilaginous fish. **a**, A conserved regulatory element downstream of Pou3f3 genes. ATAC-seq data from (top) zebrafish FACS-purified *fli1a:*EGFP; *sox10*:DsRed double-positive arch NCCs and double-negative non-arch cells, (middle) human in vitro-derived neural crest-like cells (data from ^14^), and (bottom) mouse hyoid arch cells versus early neural crest progenitors (data from ^15^). Data were visualized with the UCSC Genome Browser, and analyzed peaks are indicated. **b**, mVista alignment^28, 29^ of conserved 1B sequences relative to Hs1B. **c**, Structure of reporter construct. **d**, Compared with a *pou3f3b* knockin line (left column), Dr1 and 1B reporter lines show primarily arch-restricted GFP expression at 36 hpf (lateral perspective; arches are numbered), with an additional domain of expression in the anterior head (arrowheads) as indicated. **e**, **e’,** At 6-7 dpf (ventral perspective), Dr1 and bony vertebrate 1B sequences drive robust expression in the hyoid opercular mesenchyme (Op), with nominal expression in the more posterior arches, while cartilaginous fish 1B reporters induce expression evenly across the hyoid and posterior gill-bearing arches. Expression expands into the ventral operculum in the human and gar reporters (dotted line). mx = maxillary expression. Images in **e’** are cropped subsets highlighting the posterior arches. *sox10:*DsRed labels neural crest-derived and otic cells at 36 hpf and chondrocytes at 7 dpf. Scale bars = 50 µm.

We next examined the ability of 1B sequences from diverse vertebrates to drive arch expression patterns corresponding to their gill cover patterns. To do so, we cloned 1B sequences from three bony vertebrates (zebrafish, spotted gar, human) and two cartilaginous fishes (little skate and elephant fish) and tested their abilities to drive GFP expression in zebrafish stable transgenesis assays. At 36 hpf, 1B sequences from all five species directed GFP expression in hyoid and mandibular arch mesenchyme, with few or no GFP^+^ cells in the posterior arches (Fig. 2d). However, these patterns conspicuously diverged by 7 dpf. Whereas the zebrafish, gar, and human enhancers drove expression primarily in the operculum, skate and elephant fish enhancers drove strong expression in the operculum and all posterior gill-bearing arches (Fig. 2e). The zebrafish but not gar or human enhancers also recapitulated the clear restriction of GFP expression to the dorsal half of the operculum, as seen in the knockin line, suggesting divergence within bony fishes of the sequences conferring ventral repression. These experiments demonstrate that small shifts in the 1B enhancer, likely occurring after bony and cartilaginous fish diverged ∼430 mya^17^, can explain, at least in part, the unique arch expression patterns of vertebrate Pou3f3 homologs.

Given the strong correlation of Pou3f3 expression with gill cover pattern, we next assessed the requirement for Pou3f3 in gill cover development. We generated zebrafish *pou3f3a^el489^* and *pou3f3b^el502^* frame-shift alleles with TALENs and confirmed total loss of protein expression in double-mutants with a pan-Pou3f3 antibody (Extended Data Fig. 7a-b). The fan-shaped opercle and adjacent subopercle intramembranous bones form the primary skeleton of the operculum in teleost fishes (Fig. 3a). Whereas adult *pou3f3a* mutants appear normal, adult *pou3f3b* mutants have variably truncated gill covers (Fig. 3b) associated with reduced opercles and missing subopercle bones (Fig. 3c), as well as fragmented infraorbital bones (not shown). The opercle is entirely absent in compound *pou3f3b* mutants lacking one or both copies of *pou3f3a* (Fig. 3d), which develop cardiac edema and die at approximately 6-7 dpf. By contrast, the branchiostegal ray bone, derived from more ventral hyoid arch cells that do not express *pou3f3a/b*^18^, develops normally in double-mutants, as assessed by a *RUNX2*:mCherry osteoblast reporter line and endogenous *runx2b* expression at 3 dpf (Fig. 3e; Extended Data Fig. 7c), despite a failure of mineralization that is likely secondary to edema. Double mutants also display reductions in the hyomandibular cartilage (to which the opercle bone attaches) and its *sox9a*^+^ progenitors (Fig. 3d-f), as well as loss of the specialized joint cells (*trps1:*GFP^+^) that connect the opercle to the hyomandibula and disorganization of opercular muscles (Extended Data Fig. 7d-e). A requirement for Pou3f3 in bone and cartilage development in the dorsal (i.e. proximal) mandibular and hyoid arches is also conserved in mammals, as *Pou3f3^-/-^* mice lack squamosal and jugal bones (mandibular derivatives) and display an abnormal stapes cartilage (hyoid derivative)^10^.

**Fig. 3.**
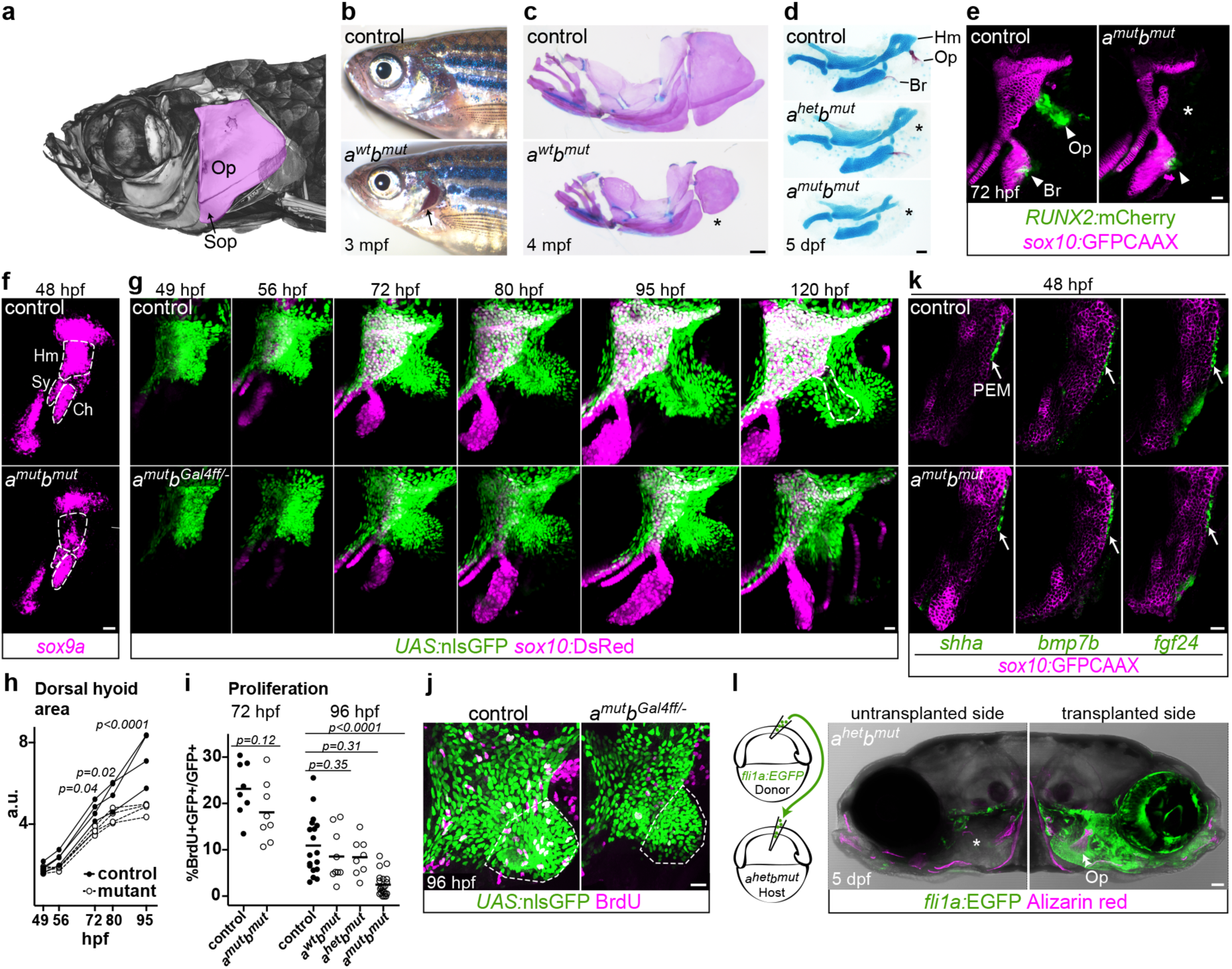
Pou3f3 is required for formation of the opercular skeleton in zebrafish. **a**, MicroCT of an adult zebrafish skull, with the opercle (Op) and subopercle (Sop) bones pseudo-colored pink. **b-c**, Reduction or loss (*) of Op and Sop bones exposes the gills (black arrow) in adult *pou3f3b* mutants. In **c**, dissected jaw skeletons were stained with Alcian blue (cartilage) and Alizarin red (bone), and the ceratohyal and brachiostegal ray series were removed for clarity. **d**, In larval mutants, the Op and supporting hyomandibula cartilage (Hm) are progressively reduced (*) with decreasing Pou3f3 dosage. **e**, Loss of the Op bone in double mutants is preceded by loss of *RUNX2:*mCherry^+^ pre-osteoblasts in this domain (*) but not the branchiostegal ray (Br) bone domain at 3 dpf. **f**, Double mutants show a reduction of *sox9a* expression in the Hm pre-cartilage condensation but not the neighboring symplectic (Sy) or ceratohyal (Ch) domains. **g**, Opercular outgrowth initiates but is not sustained in *pou3f3a; pou3f3b* mutants. Control *pou3f3a^+/-^; pou3f3b^Gal4ff/+^* and mutant *pou3f3a^-/-^; pou3f3b^Gal4ff/-^* larvae carrying *UAS*:nlsGFP and *sox10:*DsRed (cartilage) transgenes were repeatedly imaged between 49 and 120 hpf. Compaction of mesenchyme around the forming opercle bone (outlined in upper right panel) was not evident in mutants. **h**, Quantification of total dorsal hyoid arch area in individually tracked control and *pou3f3a^-/-^; pou3f3b^Gal4ff/-^* siblings (repeated measures ANOVA: genotype p=0.0069; time: p<0.0001; genotype x time: p<0.0001). **i**, A trend towards moderately lower rates of proliferation in double mutants at 72 hpf becomes significant at 96 hpf (unpaired t-test). **j**, Representative BrdU-labeled control and mutant samples, with the quantified opercular region outlined. **k**, PEM markers *shha*, *bmp7b* and *fgf24* are expressed at normal levels in *pou3f3a; pou3f3b* double mutants at 48 hpf (white arrows). *sox10*:GFPCAAX labels arch mesenchyme. **l**, Unilateral transplantation of *fli1a:EGFP* donor neural crest cells into a *pou3f3a^+/-^; pou3f3b^-/-^* host rescued Op formation. Images in **e-g**, **j**, **l** are maximum intensity projections; single optical sections are presented in **k**. Scale bars: **c** = 500 μm; **d**, **l** = 50 μm, **e-g**, **j-k** = 20 μm.

To interrogate the cellular basis of gill cover defects in Pou3f3 mutants, we analyzed *pou3f3a^-/-^; pou3f3b^Gal4ff/el502^*; *UAS:nlsGFP* embryos. As the knock-in disrupts the *pou3f3b* gene, these display similar opercular defects as *pou3f3a^-/-^*; *pou3f3b^el502/el502^* mutants (Extended Data Fig. 4a,f). Sequential confocal imaging in controls revealed posterior-directed migration of *pou3f3b*-labeled mesenchymal cells in the hyoid arch, which contributed to both *sox10*:DsRed^+^ chondrocytes of the hyomandibular cartilage and *RUNX2*:mCherry^+^ osteoblasts of the opercle bone (Fig. 3g, Extended Data Fig. 4d-e; Movie 1). In double mutants, comparable amounts of *pou3f3b*-labeled mesenchymal cells were present at 49 hpf, yet outgrowth slowed by 72 hpf and completely stalled by 95 hpf, reflected by a significant decrease in the area of the *pou3f3b*-labeled dorsal hyoid arch as early as 72 hpf (Fig. 3g-h). This failure of opercular outgrowth may be due in part to reduced proliferation, as there was a trend toward fewer BrdU^+^ *pou3f3b*-labeled cells at 72 hpf that became highly significant by 96 hpf (Fig. 3i-j; confirmed by pHH3 and PCNA staining (Extended Data Fig. 8)). The failure of sustained opercular growth is likely due to defects in the mesenchyme and not the PEM. Expression of PEM markers *shha*, *bmp7b*, and *fgf24* was unaltered in mutants at 48 hpf (Fig. 3k). Further, transplantation of wild-type neural crest precursor cells into *pou3f3a^+/-^*; *pou3f3b^-/-^* mutants fully rescued opercle bone formation, indicating that Pou3f3 function is sufficient in arch mesenchyme for development of the gill cover skeleton (Fig. 3l).

Next, we tested whether forced expression of Pou3f3b in the posterior arches was sufficient to induce the formation of ectopic gill cover-like structures in zebrafish. To do so, we used a *fli1a:*Gal4VP16 transgene to drive a *UAS*:pou3f3b transgene throughout post-migratory arch mesenchyme. At 4 dpf, Pou3f3b misexpression resulted in ectopic posterior-directed cartilaginous processes extending from the dorsal tips of the third and fourth arch-derived ceratobranchial (CB) cartilages, consistent with ectopic expression of *pou3f3b* observed in the third and fourth arches at 36 hpf (Fig. 4a,c). The facial skeleton, in particular the jaws, was also hypoplastic, potentially reflecting that *pou3f3b* was also ectopically expressed in the normally Pou3f3-negative ventral jaw-forming region. We also observed similar ectopic cartilaginous processes form in zebrafish *kat6a* (*moz*) mutants, in which Hox genes are downregulated in the posterior gill-forming arches^19^ and *pou3f3b* becomes ectopically expressed in the third and fourth arches (Fig. 4b,e). Intriguingly, the 1B enhancer is the only sequence in the murine *Pou3f3* locus bound by HOXA2 and its Pbx/Meis cofactors in ChIP-seq assays of hyoid arch cells^20^ (Fig. 2a), suggesting that Hox transcription factors might act through the 1B enhancer to prevent posterior arch expression of Pou3f3 orthologs in the bony fish clade. Whereas posterior arch expression of *pou3f3b* was linked to ectopic skeletal elements, lethality by 7 dpf in both *pou3f3b*-misexpression and *kat6a* mutant embryos precluded us from assessing whether more extensive ectopic gill covers might form. We also noted no ectopic *shha^+^* PEMs in gill arch epithelia of *pou3f3b*-misexpression or *kat6a^-/-^* embryos (Fig. 4d,f)^19^. Generation of the full multiple gill cover state seen in cartilaginous fishes might therefore require the establishment of PEMs in posterior arch epithelia in addition to expression of Pou3f3 orthologs in posterior arch mesenchyme.

**Fig. 4.**
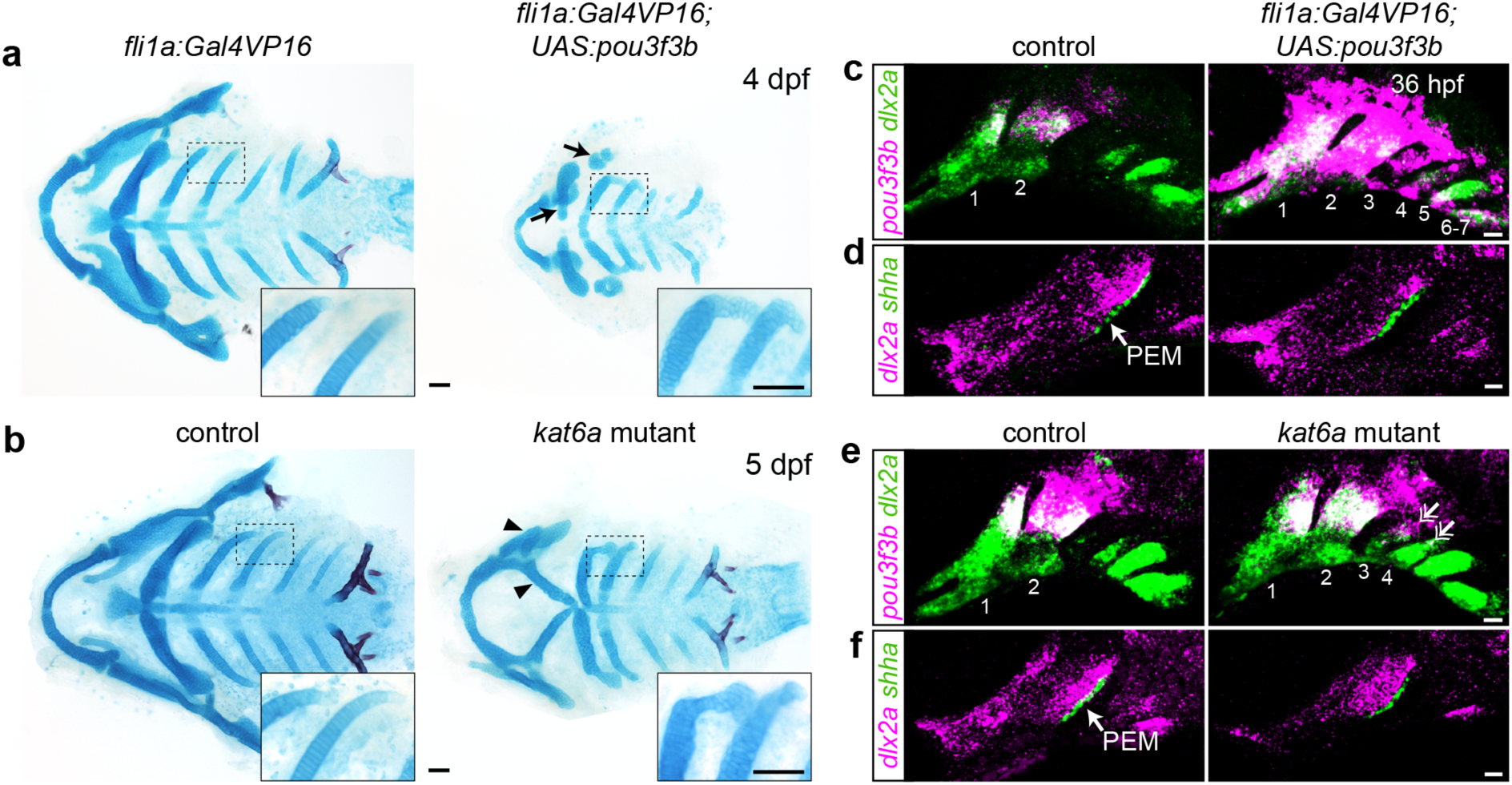
Misexpression of *pou3f3b* in posterior arch mesenchyme induces ectopic posterior-directed cartilages. Cartilage (blue) and bone (red) staining reveals ectopic cartilaginous projections in the third and fourth arches (insets) of *fli1a:Gal4VP16*; *UAS:pou3f3b* (**a**) and *kat6a^-/-^* (**b**) larvae. The hyoid skeleton was reduced in *fli1a:Gal4VP16*; *UAS:pou3f3b* embryos (arrows in **a**) and homeotically transformed in *kat6a* mutants (arrowheads in **b**). **c**, **e**, In situ hybridization for *pou3f3b* (pink) relative to the pan-mesenchyme marker *dlx2a* (green) shows ectopic expression of *pou3f3b* in neural crest-derived mesenchyme (primarily arches 1-4) in *fli1a:Gal4VP16*; *UAS:pou3f3b* embryos and in the dorsal third and fourth arches (double arrows) of *kat6a* mutants. **d, f**, In situ hybridizations show that the PEM marker *shha* remains confined to the hyoid arch (arrow) in *fli1a:Gal4VP16*; *UAS:pou3f3b* and *kat6a^-/-^* embryos. In situ images are maximum intensity projections. Scale bars: **a**, **b** = 50 μm; **c-f** = 20 μm.

We propose that in an ancestral gnathostome, co-option of the neural gene Pou3f3 into the arches, through acquisition of the 1B arch enhancer, conferred a new capacity for mesenchymal growth and skeletal differentiation that led to the formation of a gill cover. Recent discoveries of a stem gnathostome^21^ and a stem chondrichthyan^22^ with prominent hyoid arch bony opercula are consistent with a scenario in which the last common ancestor of extant jawed vertebrates possessed a single, hyoid arch-derived gill cover. The four posterior gill covers of cartilaginous fishes would therefore have evolved after the two lineages diverged^23^, perhaps coincident with sequence changes in the 1B enhancer permitting robust *Pou3f3* expression in the posterior arches. Sequence modifications in otherwise highly conserved enhancers have also been linked to other more recent evolutionary transitions, e.g., the loss of limbs in snakes (Shh ZRS enhancer^24^) and the fin-to-limb transformation in tetrapods (Gli3 intronic enhancer^25^). It remains to be determined whether other opercular novelties are also associated with modifications to the 1B sequence or Pou3f3 activity more generally, such as the precocious opercular outgrowth with external gill development observed in bichir ^26^ or the disappearance of the entire opercular skeleton in ancestral tetrapods upon their migration onto land^2, 27^.

## Movie 1. Posterior-directed growth of *pou3f3b*-expressing hyoid mesenchyme

Timelapse imaging of a control *pou3f3b^Gal4ff^; UAS:*nlsGFP; *sox10*:DsRed embryo from 56 to ∼72 hpf. Note the posterior migration of GFP^+^ hyoid cells to form the opercular flap. Anterior is to the left. *sox10:*DsRed (magenta) is expressed by chondrocytes in the facial skeleton (middle) and the pectoral fin (top right) as well as the otic vesicle (top center).

## Methods

The Institutional Animal Care and Use Committees of the University of Southern California (No. 10885, 20540), Cincinnati Children’s Hospital Medical Center (No. 2018-0076), the University of Colorado at Boulder (No. 2392), and the Marine Biological Laboratory (No. 17-31, 18-32) approved all of the animal procedures carried out in this study. Mice were housed in accordance with National Institutes of Health Guidelines.

### Husbandry and specimen collection

Zebrafish (*Danio rerio*) were reared in embryo medium^30^ at 28.5°C and staged according to ^31^. Mixed-background wild-type mouse embryos were collected from timed-pregnant females at 9.5 and 11.5 days post coitus. Adult sea lamprey (*Petromyzon marinus*) were maintained and embryos were raised and collected as described^32^. Skate embryos (*Leucoraja erinacea*) were reared to relevant stages in a flow-through seawater system at 15°C at the Marine Biological Laboratory in Woods Hole, MA, and were fixed as described^33^. Holocephalan embryos (*Callorhinchus milii*) were collected from the field and fixed as described^3^.

### Zebrafish lines and genotyping

Existing mutant and transgenic lines used in this study include *kat6a*/*moz^b719^* 19, *trps^^<u>j1271aG</u>t^* (referred to as *trps1:GFP*)^34^, *Tg(fli1a:Gal4VP16)^el360^* ^35^, *Tg(fli1a:eGFP)^y1^* ^36^, *TgBAC(hand2:eGFP)^pd24Tg^* ^37^, *Tg(RUNX2:mCherry)* (gift of Shannon Fisher), *Tg(sox10:GFPCAAX)* ^38^, and *Tg(sox10:DsRedExpress)^el10^* ^39^. Two new mutant lines (*pou3f3a^el489^* and *pou3f3b^el502^*), a targeted knock-in line (*pou3f3b^Gal4ff-el795^*), and two transgenic lines (*Tg(UAS:pou3f3b; α-crystallin:Cerulean)^el578^* and *Tg(UAS:nlsGFP; α-crystallin:Cerulean)^el609^*) were generated as part of this study (see below). Transgenic lines were maintained by selectively raising larvae expressing fluorescent marker proteins. To identify carriers of the mutant alleles, the caudal fin was biopsied under tricaine anesthesia (Western Chemicals) at 14 or 90 dpf, and the tissue was digested with proteinase K and genotyped by PCR/restriction fragment length polymorphism assays using GoTaq DNA polymerase (Promega). The primers used to genotype the *pou3f3a* and *pou3f3b* mutant lines were pou3f3a-F2 5’-ACCACGCATACTTTTCCAGC-3’ and pou3f3a-R2 5’-CTCCTTGCATGAAGTCGCTC-3’; pou3f3b-Fout 5’-TCGATAGTGCACTCGGACTC-3’ and pou3f3b-Rout 5’-CCAGGCTGCGAGTATATGAGA-3’. The reaction was run at 95°C for 3 min, followed by 35 cycles of 95°C for 15 s, 58°C for 30 s, and 72°C for 30 s, with a final extension at 72°C for 5 min. The 311- and 473-bp products for *pou3f3a* and *pou3f3b* were digested with HinfI at 37°C and ApaI at 25°C, respectively, which cut the wild-type alleles only.

### Generation of new zebrafish mutant and transgenic lines

*pou3f3a* and *pou3f3b* mutant lines were created with TALENs targeting the following sequences: pou3f3a-L: 5’-TCTCTCATCAGCCTCGCTCG-3’ and pou3f3a-R 5’-TCCCGGCTGCATGCCACCAC-3’; pou3f3b-L: 5’-TGCACTCTGGGACTGCGCTG-3’ and pou3f3b-R: 5’-TCTGGTGGGGACCTAAATGT-3’. TALEN constructs were assembled using a PCR-based platform^40^, digested with StuI (New England Biolabs, Ipswich, MA), and used as templates for RNA synthesis with the mMessage mMachine T7 Ultra kit (Ambion/Life Technologies, Carlsbad, CA, USA). Pairs of TALEN RNAs were injected into 1**-**cell-stage embryos at 100 ng/μl, and mosaic germline founders were identified by screening their offspring for frameshift alleles using PCR and restriction digest assays followed by Sanger sequencing. The *pou3f3a^el489^* allele is an 8-bp deletion that causes a frameshift after amino acid 23 (of 438), leading to a premature stop codon (PTC) after 34 incorrect amino acids. *pou3f3b^el502^* is a 2-bp deletion that results in a frameshift after amino acid 135 (of 425) and a PTC after 15 incorrect amino acids. Both mutations occur upstream of the conserved POU and homeobox domains. No *pou3f3^+/-^; pou3f3b^-/-^* or double mutants were recovered as adults, though single *pou3f3a* and *pou3f3b* mutants and *pou3f3a^-/-^; pou3f3b^+/-^* fish were fully viable and fertile.

The *pou3f3b^Gal4ff-el795^* targeted knockin line was made using a CRISPR/Cas9-based protocol^41^. Briefly, three guide RNAs targeting sequences upstream of the *pou3f3b* translational start site (5’-AAACATATTCATAAGGTTAA-3’, 5’-GGTTAACGGAATGGCCACAG-3’, and 5’-AGCAAAGAGAAAGTATCTGC-3’) were co-injected at 100 ng/µl into *Tg(UAS:nls:GFP)* embryos together with Cas9 RNA (100 ng/µl), a circular *hsp70l:Gal4ff:pA* construct, and a fourth gRNA targeting a bait sequence within the construct, which linearized the donor DNA *in vivo*^41^. Germline founders that recapitulated the embryonic expression pattern of *pou3f3b* were identified by crossing back to *Tg(UAS:nls:GFP)* upon reaching maturity.

The UAS:Pou3f3b; α-crystallin:Cerulean and UAS:nlsGFP; α-crystallin:Cerulean constructs were created with the Gateway Tol2kit^42^ (Invitrogen) by combining pMEs containing the *pou3f3b* or *nlsGFP* coding sequence with p5E-UAS, p3E-polyA and pDestTol2AB2 in an LR reaction. The stable lines *Tg(UAS:pou3f3b; α-crystallin:Cerulean)el578* and *Tg(UAS:nlsGFP; α-crystallin:Cerulean)el609* were established by injecting each construct together with transposase RNA (30 ng/µl each) into wild-type embryos and selecting for fish whose offspring expressed Cerulean in the lens.

To test the function of putative Pou3f3 enhancers, sequences downstream of *pou3f3a* or *pou3f3b* that were enriched for open chromatin in *fli1a:E*GFP^+^; *sox10:*DsRed^+^ vs. *fli1a:E*GFP^-^; *sox10:*DsRed^-^ cells (Dr4, Dr1, Dr1A, and Dr1B) were PCR-amplified from genomic DNA and cloned into pDONRp4p1r to generate p5E Gateway vectors. gBlocks were generated for the human, spotted gar, little skate, and elephant fish 1B homologs (Integrated DNA Technologies; Extended Data Table 1) and used to make p5E vectors. All p5Es were combined with pME-E1B:GFP, p3E:polyA, and pDestTol2AB2 to generate reporter constructs that use the minimal E1B promoter to drive GFP expression and carry the *α-crystallin*:Cerulean marker. These were microinjected with transposase as described above. Two independent stable alleles per construct were analyzed in the F1 and/or F2 generation.

### In situ hybridization

Published zebrafish probes include *dlx2a*^43^, *pou3f3a, pou3f3b-3’UTR*^9^, *runx2b*^44^, *shha*^45^, and *sox9a*^46^. cDNAs for an exon-only *pou3f3b* probe, *bmp7b*, *fgf24*, lamprey *Pou3a*, and mouse *Pou3f3* were PCR-amplified by Herculase II Fusion DNA Polymerase (Agilent) (Extended Data Table 2) and cloned into pCR-Blunt II-TOPO or pJet1.2 (Thermo Fisher Scientific, Waltham, MA). Fragments of skate and holocephalan *Pou3f3* were PCR-amplified from total embryonic cDNA using REDTaq DNA polymerase (Millipore Sigma, St. Louis, MO) and cloned into the pGemT-easy vector system (Promega, Madison, WI). Lamprey *Pou3b* and *Pou3c* cDNA sequences were purchased as fragments cloned into pUC57 (Synbio Technologies, Monmouth Junction, NJ) (Extended Data Table 3), and RNA probe templates were PCR-amplified with M13 primers. In all other cases, plasmids were linearized, and antisense probes incorporating dioxygenin (DIG)- or dinitrophenol (DNP)-labeled nucleotides were synthesized with Sp6 or T7 polymerase (Promega, Roche).

Embryos were collected at the designated stages and fixed overnight in 4% paraformaldehyde (PFA), passed through a methanol gradient and stored at −80°C until use. In situ hybridizations were performed as previously described for zebrafish^47^, mouse^48^ (modified to include maleic acid buffers), skate, elephant fish ^49, 50^ and lamprey^51^. A minimum of three mutant or transgenic embryos was compared to a similar number of control siblings, and representative images are presented.

### Staining and cell proliferation assays

The cranial musculature was labeled by incubating PFA-fixed *sox10:GFPCAAX^+^*; *pou3f3a; pou3f3b* mutant and control larvae with Alexa633-conjugated phalloidin (1:100 in PBS) following permeabilization in five 60-min washes of 1% Triton-X-100. Alcian Blue and Alizarin Red staining to reveal cartilage and bone structure was performed as previously described for larvae^52^ and juvenile/adult fish^53^. For immunostaining, PFA-fixed embryos or larvae were passed through a methanol gradient and permeabilized with cold acetone at −20°C for 7-13 min prior to immunostaining with anti-Pou3f3 (1:200, Abcam ab180094), anti-pHH3 (1:500, Millipore Sigma 06-570), or anti-PCNA (1:150, Thermo Fisher 13-3900) and anti-GFP (1:200, Abcam ab13970) primary antibodies and Alexa568- and Alexa488-conjugated secondary antibodies.

To acquire a snapshot of proliferating cells in the growing operculum, 72 and 96 hpf larvae were incubated with 4.5 mg/ml BrdU in 15% DMSO for 10 min, then fixed in 4% PFA for 3 h at room temperature, passed through a methanol gradient and stored at −80°C. Larvae were subsequently rehydrated, genotyped, digested with proteinase K (20 µg/ml) at 25°C for 5 min, post-fixed in 4% PFA for 20 min, permeabilized in cold acetone for 13 min, then treated with 2N HCl for 1 h at 25°C before immunostaining with anti-GFP (1:300) and anti-BrdU (1:200, Novus Biologicals NB500-169) primary antibodies and Alexa568- and Alexa488-conjugated secondary antibodies in blocking solution containing 0.5% Triton-X-100.

### Transplants

Unilateral cell transplants were performed as described^54^, with the contralateral side serving as an internal control. In brief, naïve ectoderm from *fli1a:EGFP* donor embryos was transplanted into the neural crest fate-mapped domain of *pou3f3a; pou3f3b* mutant hosts at 6 hpf. At 5 dpf, transplanted larvae were live-stained with 140 µM Alizarin red for 30 min prior to imaging, and host genotype was confirmed by tail biopsy.

### Imaging

Colorimetric in situs and adult skeletons were imaged using LAS software with a Leica S8APO or M165FC stereomicroscope, and larval skeletons with a Leica DM2500 compound microscope. Adult *pou3f3b* fish were imaged immediately following euthanasia with an AxioCam MRm on a Zeiss StereoDiscovery microscope. Transgenic or fluorescently stained samples were imaged with a Zeiss LSM800 or a Nikon A1R inverted confocal and are shown as single optical sections or maximum intensity projections, as indicated. Brightness and contrast were modified evenly across samples using Adobe Photoshop CS6 or CC2019. Time-lapse imaging of control *pou3f3b^Gal4ff^; UAS:*nlsGFP*; sox10:*DsRed fish was performed with a 20x objective beginning at 56 hpf as previously described^54^, with a 61.2 µm z-stack imaged every 18 minutes.

### Data analysis

The numbers of BrdU^+^GFP^+^, PCNA^+^GFP^+^ or pHH3^+^/GFP^+^ cells within the operculum were determined using the Spot Colocalization MATLAB extension in Imaris (Bitplane). Genotypes (n > 6 each) were compared using unpaired two-tailed t-tests (assuming normal distributions), with p < 0.05 deemed significant. Area of the dorsal hyoid arch was calculated using the polygon tool in ImageJ (NIH) and analyzed by two-way repeated measures ANOVA (alpha = 0.05) followed by Bonferroni’s test for multiple comparisons. Statistical tests were performed using GraphPad Prism v8.

### ATAC-seq

*fli1a:EGFP*; *sox10:DsRed* double-positive embryos were selected under a fluorescent dissecting microscope at 36 hpf, dechorionated, and dissociated in batches of 25 as previously described^55^. Cells were sorted based on GFP and DsRed expression on a MoFlo Astrios instrument (Beckman-Coulter, Brea, CA). Approximately 15,000 double-positive cells and 20,000 double-negative cells were collected into PBS with 5% FBS and used to construct ATAC-seq libraries using a low-input μATAC-seq protocol (manuscript in preparation, modified from ^56^). Briefly, cells were centrifuged at 700 x g, and the supernatant was carefully removed using a 200-μl pipette tip with a broad opening. Next, 20 μl lysis buffer (10 mM Tris-HCl pH 7.4, 5 mM MgCl_2_, 10% DMF, 0.2% NP40) was added to the cell pellet and mixed by pipetting up and down 5-10 times. A 30-μl aliquot of reaction buffer (10 mM Tris-HCl pH 7.4, 5 mM MgCl2, 10% DMF) containing homemade Tn5 transposase^57^ was then immediately added to the lysis, and the samples were incubated for 20 min at 37℃. DNA fragments were extracted with the Qiagen MinElute kit and amplified by PCR for 5 cycles using NEBNext Master Mix and indexed primers. Additional cycles were determined by performing qPCR on 1/10 of the amplified libraries. Following amplification, the libraries were cleaned using AMPURE beads to preserve only those fragments longer than 100 bp. The libraries were then pooled together before sequencing on an Illumina NextSeq 550, generating 75-bp paired-end reads that were aligned to GRCz10 with Bowtie2 and ENCODE settings.

### Data availability

ATAC-seq data are available as NCBI GEO accession GSE140636.

## Acknowledgments

We are grateful to Megan Matsutani, Jennifer DeKoeyer Crump, and Sandhya Paudel for fish care, Jeffrey Boyd at the USC Stem Cell Flow Cytometry Core Facility for FACS, and the USC Norris Medical Library Bioinformatics Service for sequence analysis. Funding was provided by the National Institute of Dental and Craniofacial Research [R35 DE027550 (J.G.C), R00 DE026239 (L.B.), R21 DE025940A (D.M.M)], the National Institute on Deafness and Other Communication Disorders [R01 DC015829 (to Neil Segil, supporting H.V.Y.)], the National Science Foundation [IOS 1744837 (D.M.M.)], the A.P. Giannini Foundation (L.B.), startup funds from Cincinnati Children’s Research Foundation (L.B.), the Scientific Grant Agency of Slovak Republic [VEGA 1/0415/17 (D.J.)], Royal Society University Research Fellowship UF130182 and Isaac Newton Trust award 14.23z (J.A.G.), and a BBSRC Doctoral Training Partnership studentship (C.H.). The bioinformatics software and computing resources were funded by the USC Office of Research, the USC Norris Medical Library, Cincinnati Children’s Research Foundation, and the Hearing Health Foundation.

## Author contributions

The project was conceived by L.B. and J.G.C. Zebrafish experiments were performed by L.B., P.F., P.X., N.N., and J.G.C; lamprey studies by D.J., T.S., and D.M.M.; and skate/holocephalan studies by C.H. and J.A.G. ATAC-seq was carried out by P.X. and H.V.Y. Writing and interpretation were performed primarily by L.B. and J.G.C. with input from P.F., D.J., D.M.M., and J.A.G.

## Competing interests declaration

The authors declare no competing interests.

## Extended Data

**Extended Data Fig. 1.**
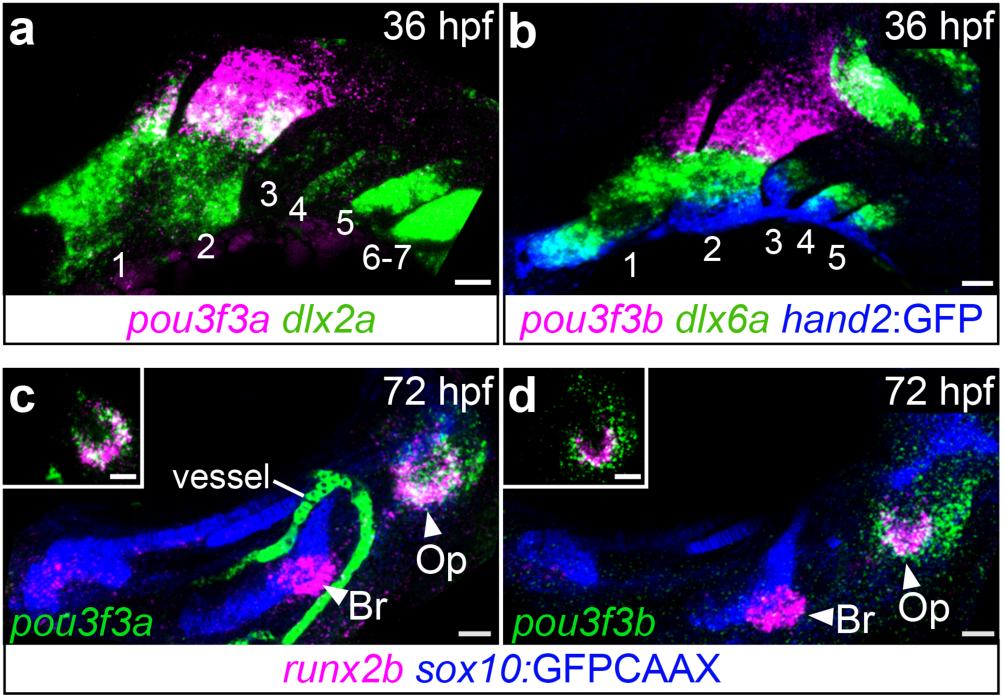
*pou3f3a/b* expression is restricted to the mandibular and hyoid arches and later to the opercle-forming domain. **a-b**, Unlike most dorsoventral patterning genes, such *dlx2a*, *dlx6a* and *hand2*, which are upregulated in each arch as they form from anterior to posterior, *pou3f3a* and *pou3f3b* remain restricted to arches 1 and 2. Visible arches are indicated in each panel. **c-d**, By 72 hpf, *pou3f3a/b* expression is further restricted to the hyoid arch-derived operculum, overlapping the semicircle of *runx2b*^+^ pre-osteoblasts at the ventral end of the forming opercle (Op) bone but not the branchiostegal ray (Br) bone. Insets in **c-d** are single optical sections; all others are maximum intensity projections. *sox10:*GFPCAAX marks the facial cartilages. Scale bars = 20 µm.

**Extended Data Fig. 2.**
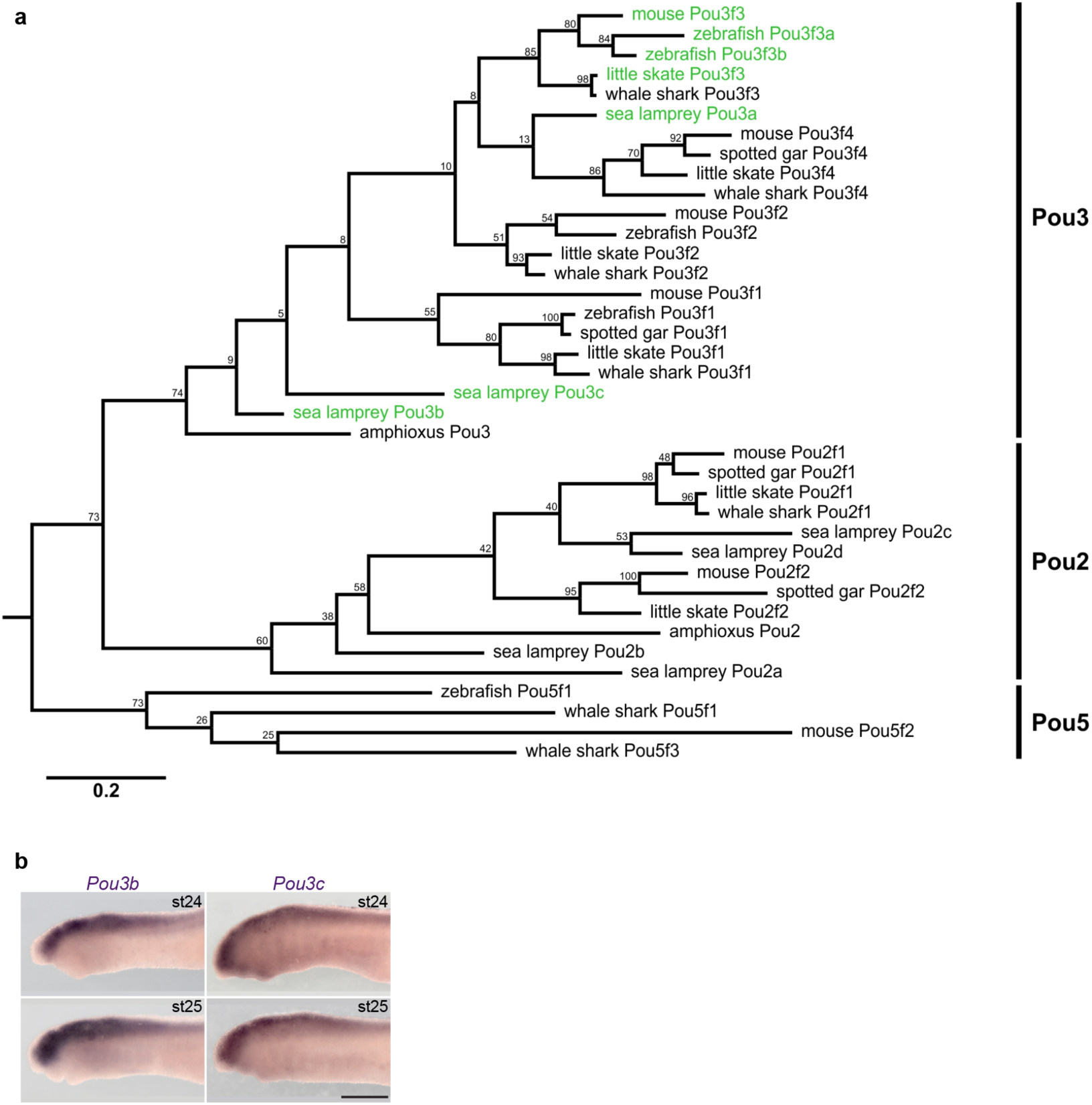
Phylogenetic analysis of POU domain homeobox genes reveals three divergent Pou3 homologs in the lamprey genome. **a**, To identify possible lamprey Pou3f3 homologs, we retrieved lamprey (*Petromyzon marinus*) Pou-domain containing sequences from a larval transcriptome and determined their similarity to gnathostome Pou sequences. Two transcript translation products clustered at the base of the gnathostome Pou3 class (termed *Pou3b* and *Pou3c*, while one showed an affinity to Pou3f4 (*Pou3a*). The results of a bootstrap analysis with 100 replications are shown at every node as the percentage of replicates that supported the branch position. Although this bootstrap analysis did not provide strong support for the specific positions of these short lamprey Pou3 fragments, these sequences are the only lamprey sequences we identified that show a specific affinity for the Pou3 group, to the exclusion of the closely related lamprey Pou2 orthologs we also identified. The *Pou3a* and *Pou3c* genes appear to be incompletely assembled in the 2017 petMar3 germline genome and are located on short contigs lacking other genes, preventing us from verifying their orthology by synteny. *Pou3b* is located on contig 54v1 near the *Klhl32* gene, a gene that is also present at the gnathostome *Pou3f2* locus. **b**, In situ hybridizations for lamprey *Pou3b* and *Pou3c* show expression in the CNS but not the pharyngeal arches, similar to *Pou3a*. Scale bar = 250 μm.

**Extended Data Fig. 3.**
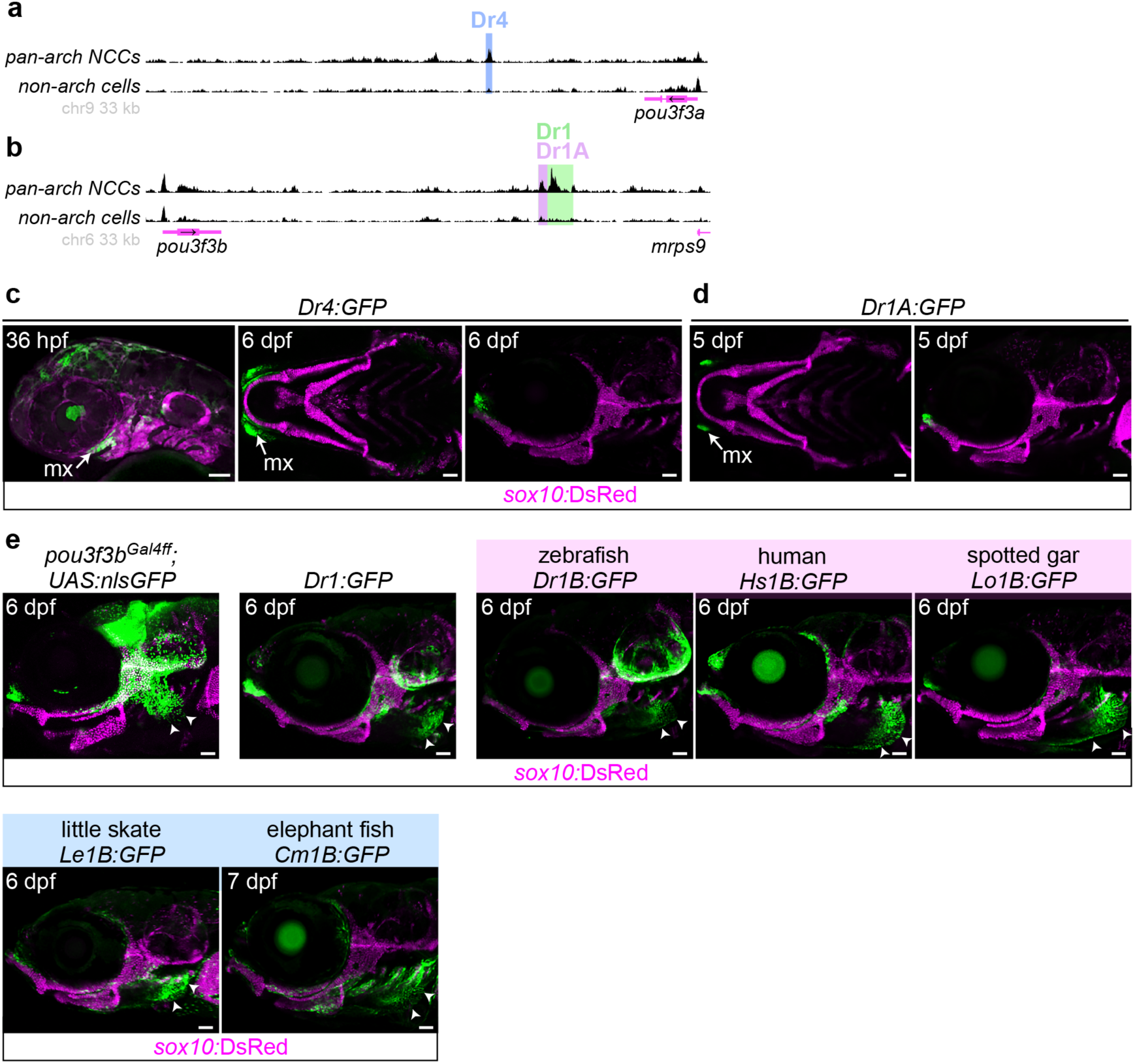
Dr4 and Dr1A enhancers do not drive strong opercular expression compared with 1B enhancers. **a-b**, The Dr4 (12.3 kb downstream of the *pou3f3a* TSS) and Dr1A (21.7 kb downstream of *pou3f3b*) enhancers were identified via enrichment for open chromatin in FACS-purified *fli1a:*EGFP; *sox10*:DsRed double-positive (arch NCCs) versus double-negative (non-arch) cells. ATAC-seq data are visualized with the UCSC Genome Browser. **c**, Transgenic reporter lines show that Dr4 drives maxillary, mandibular and hyoid expression at 36 hpf, but minimal opercular expression at 6-7 dpf. **d**, *Dr1A:*GFP is restricted to the maxillary domain at 5 dpf. **e**, Lateral views of the *pou3f3b* Gal4ff knockin line compared with the Dr1 and 1B enhancer lines at 6 dpf. Note the enrichment of GFP expression at the leading edge of the growing operculum and lack of CNS expression in all enhancer lines. Scale bars = 50 µm.

**Extended Data Fig. 4.**
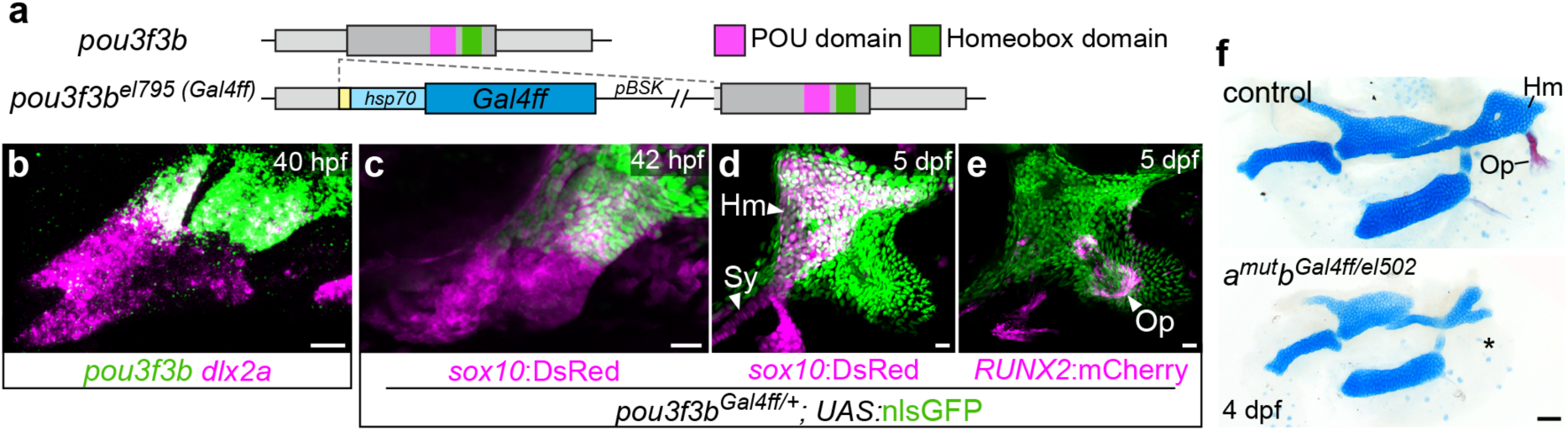
*pou3f3b* knockin line recapitulates endogenous expression and behaves as a null allele. **a**, A linearized pBluescriptSK vector containing a CRISPR target site (yellow) and the *hsp70l* minimal promoter driving Gal4ff was knocked into the *pou3f3b* 5’UTR to create allele el795 (*pou3f3b^Gal4ff^*). **b**, When crossed to a *UAS:*nlsGFP reporter, expression is observed in the CNS, periotic mesenchyme, and arches 1-2 (**b’**), mimicking endogenous *pou3f3b* mRNA expression at 40 hpf (**c**). *sox10:*DsRed and *dlx2a* label arch neural crest cells. **d**, At 5 dpf, the reporter marks the dorsal hyoid-derived hyomandibula (Hm) but not the symplectic (Sy) cartilage (*sox10:*DsRed^+^) as well as the opercular flap and opercle bone (Op, *RUNX2:*mCherry^+^) (**e**). Scale bars = 20 μm. **f**, *pou3f3a^-/-^; pou3f3b^Gal4ff/el502^* larval skeletons lack the opercle bone (asterisk) and have a forked hyomandibula, similar to *pou3f3a^-/-^; pou3f3b^el502/el502^* mutants. Scale bar = 50 µm.

**Extended Data Fig. 5.**
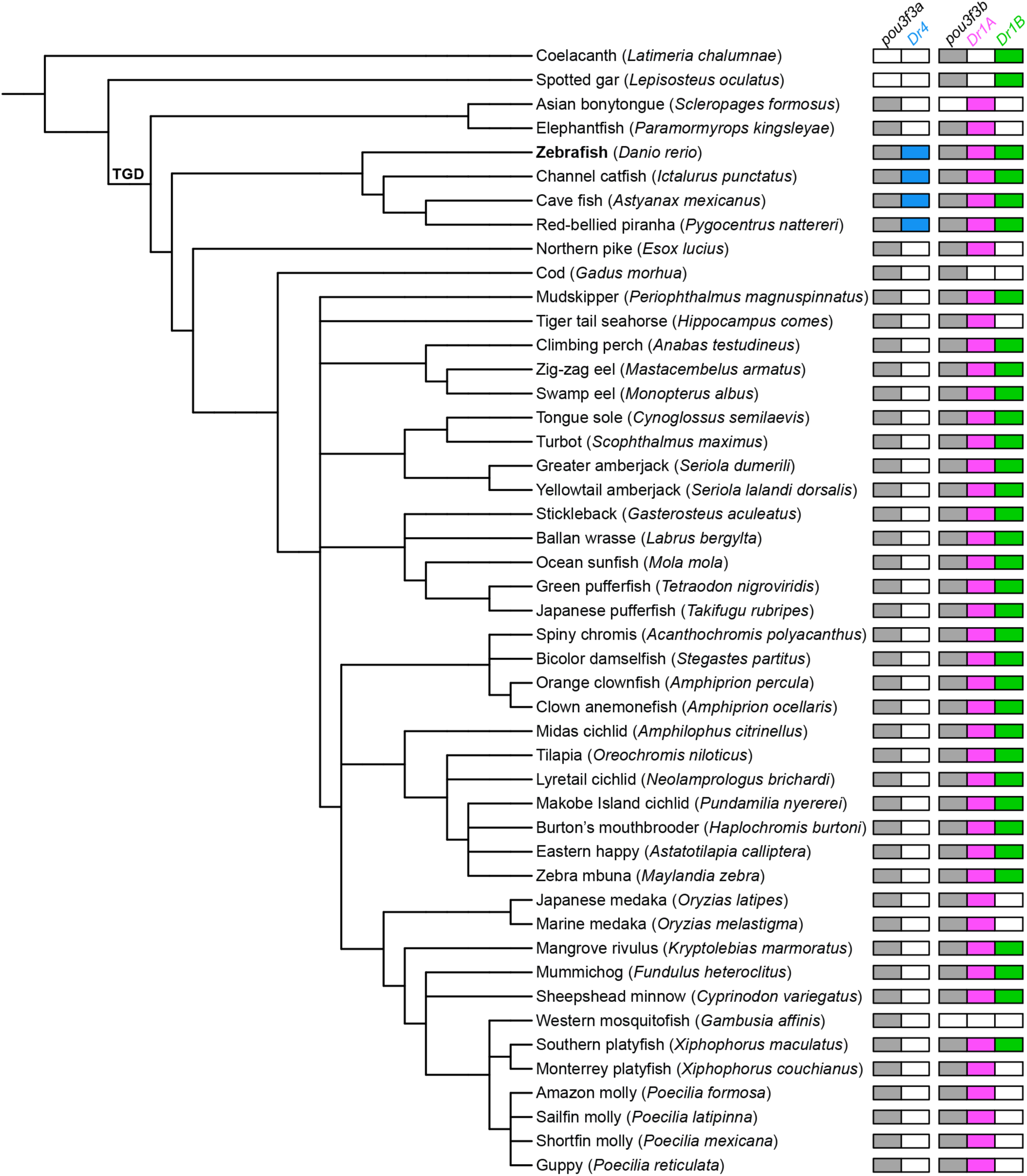
Conservation of *pou3f3* genes and associated regulatory sequences in fishes. Colored boxes next to each species name indicate that homologs of the *pou3f3a* or *pou3f3b* coding sequences or the Dr4, Dr1A, or Dr1B enhancers were successfully retrieved by BLASTing with either the zebrafish or turbot homolog. Tree generated using iTOL^58^. TGD = teleost genome duplication.

**Extended Data Fig. 6.**
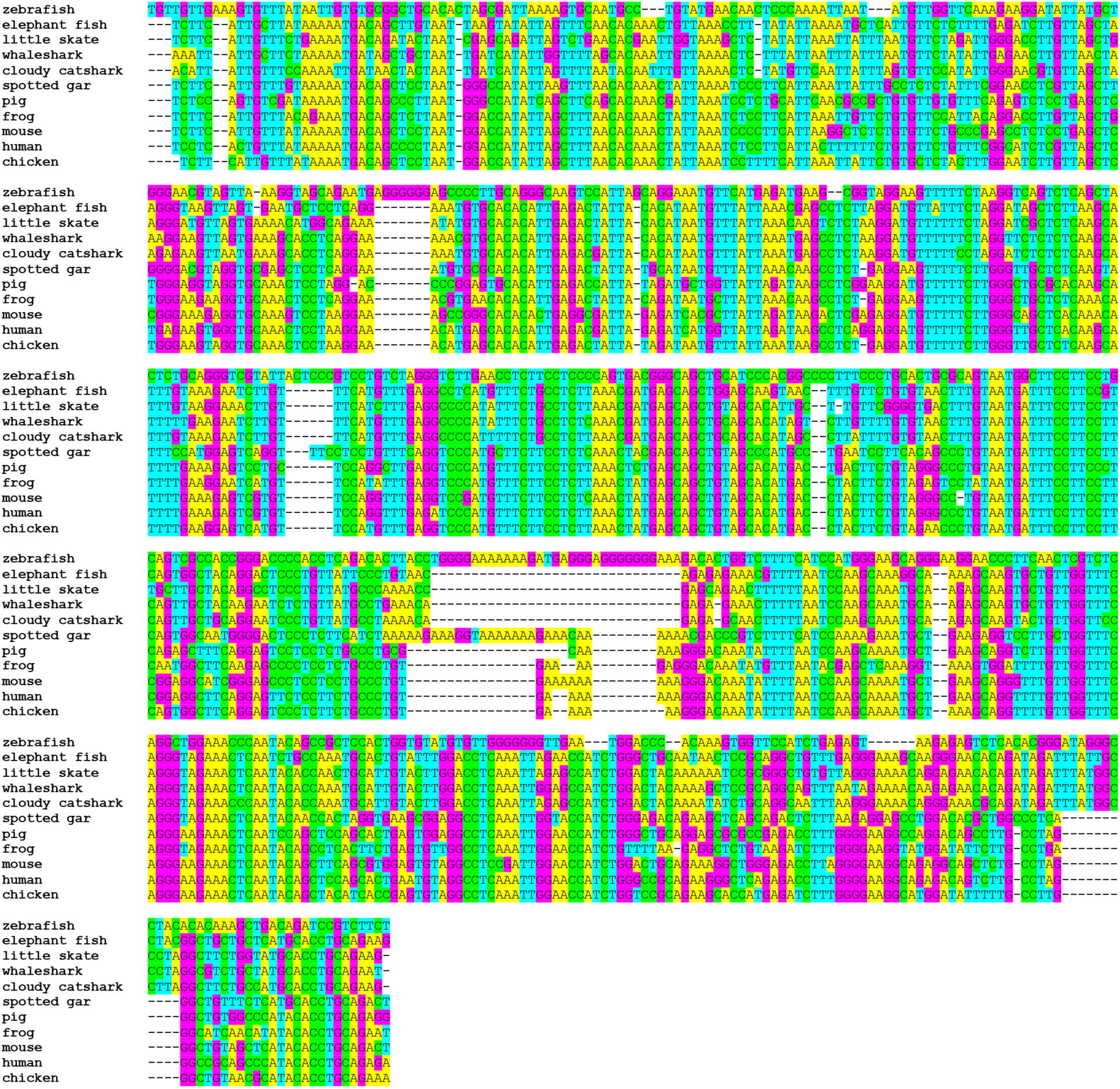
Multiple sequence alignment of eleven vertebrate *Pou3f3* 1B enhancers. 1B enhancer sequences retrieved from representative bony fish (zebrafish, spotted gar, frog, chicken, pig, mouse, human) and cartilaginous fish (little skate, cloudy catshark, whaleshark, elephant fish) genomes were aligned by Clustal Omega. Sequence identity values ranged from 51-76% for bony vs. cartilaginous species, 50-89% across bony fishes, and 82-90% across cartilaginous fishes.

**Extended Data Fig. 7.**
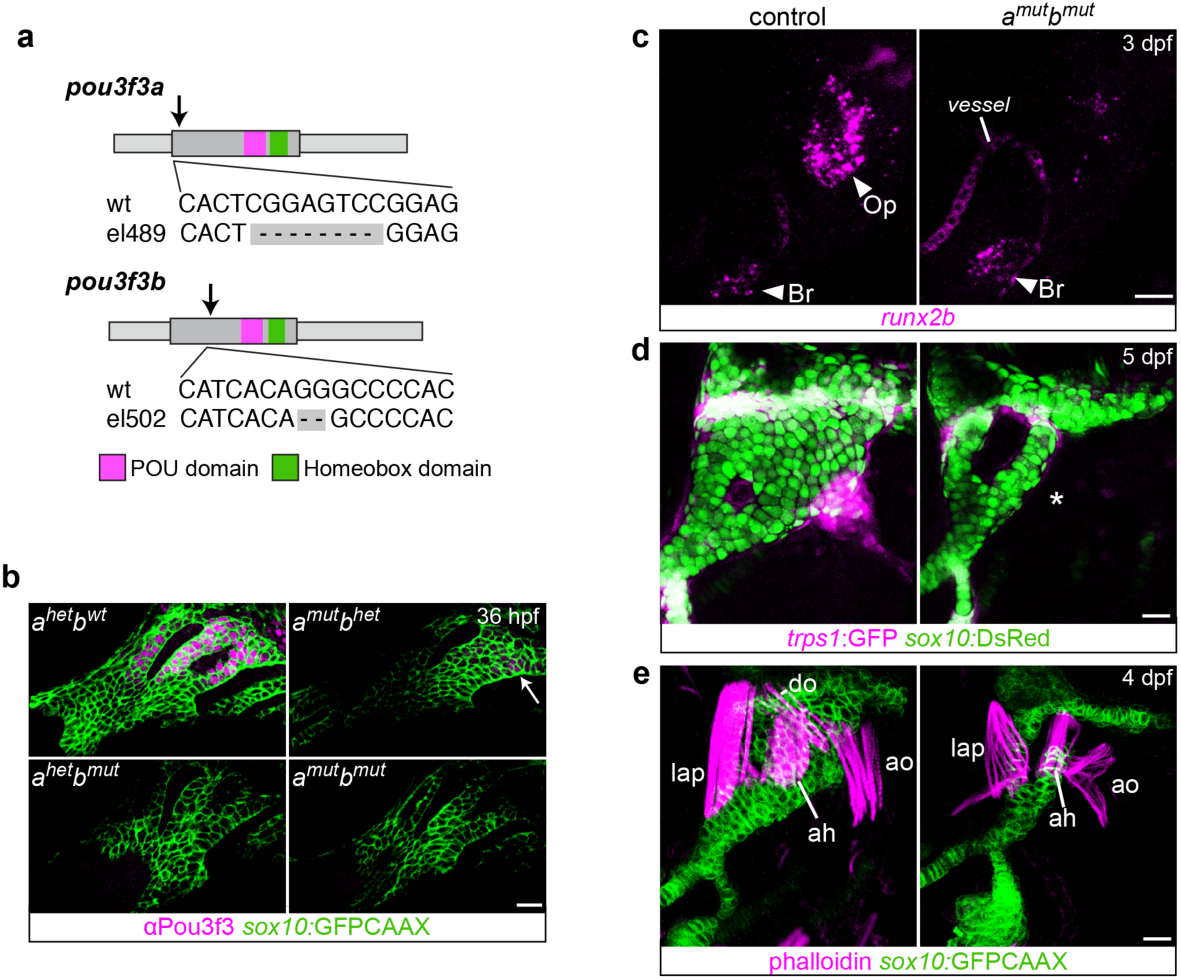
Zebrafish *pou3f3a* and *pou3f3b* TALEN mutants. **a**, Mutations were targeted to the 5’ end of each gene, upstream of the POU and homeobox domains (black arrows). **b**, Reduced pan-Pou3f3 protein expression in *pou3f3a^-/-^; pou3f3b^+/-^* embryos (white arrow), and no expression in *pou3f3a^+/-^; pou3f3b^-/-^* or double mutants at 36 hpf. The antibody recognizes the C-terminus of the protein. *sox10:*GFPCAAX labels arch NCCs. **c**, *runx2b*^+^ osteoblasts are specifically lost in the dorsal opercle (Op) but not the ventral branchiostegal ray (Br) domains in *pou3f3a^-/-^; pou3f3b^-/-^* mutants at 72 hpf. **d**, The *trps1:*GFP^+^ joint connecting the Op to the Hm cartilage does not form in the mutant (asterisk). **e**, Phalloidin staining reveals that the muscles that abduct and adduct the Op bone (dilator operculi (do) and adductor operculi (ao)) and Hm cartilage (levator arcus palatini (lap) and adductor hyoideus (ah)) are absent (do) or disorganized (ao, lap, ah) in the mutant. *sox10:*DsRed and *sox10:*GFPCAAX mark chondrocytes. Images in **b** are single optical sections; **c-e** are maximum intensity projections. Scale bars = 20 μm.

**Extended Data Fig. 8.**
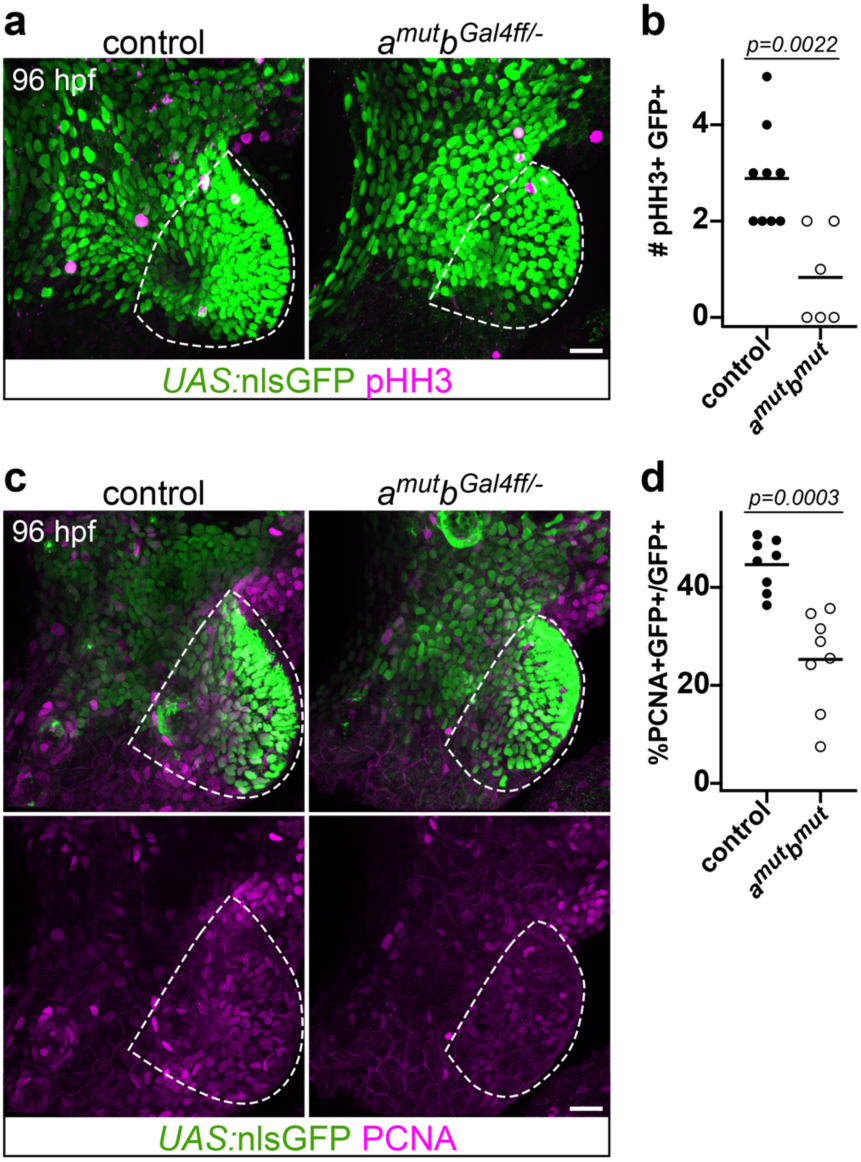
Cell proliferation is reduced in the *pou3f3* mutant operculum. Significantly fewer pHH3^+^/GFP^+^ (**a-b**) and PCNA^+^/GFP^+^ cells (**c-d**; counted relative to total GFP^+^ cells) are observed in the mutant opercular flap at 96 hpf (unpaired t-test; line indicates the mean). The quantified opercular region is outlined. Scale bars = 20 µm.

**Extended Data Table 1.**
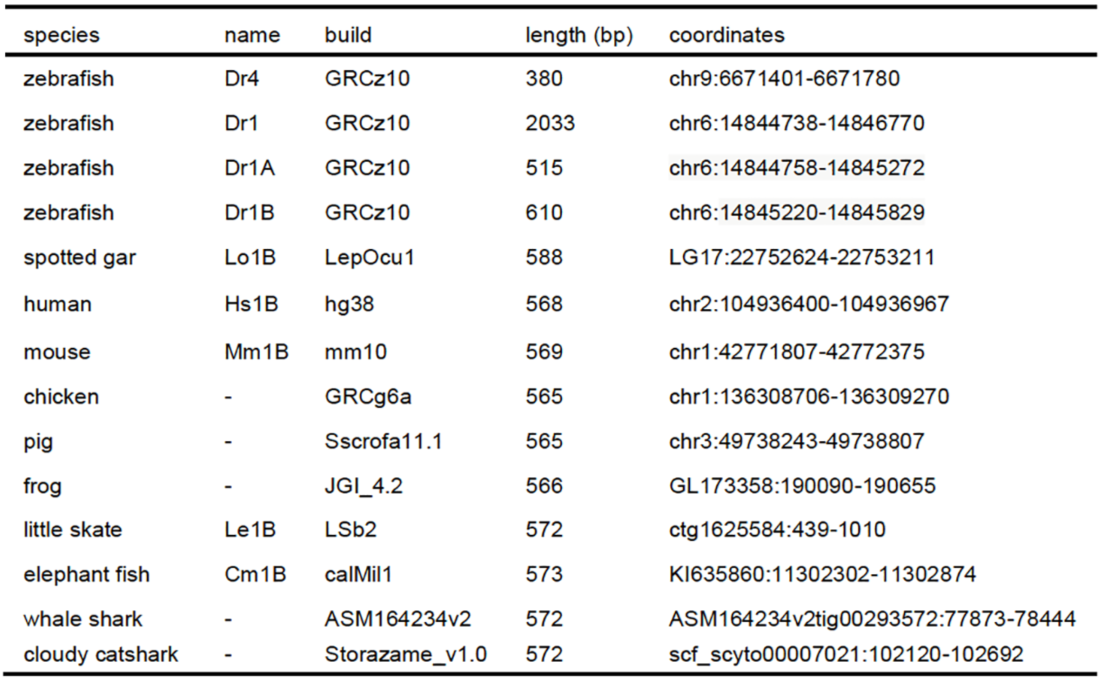
Pou3f3 enhancers.

**Extended Data Table 2.**
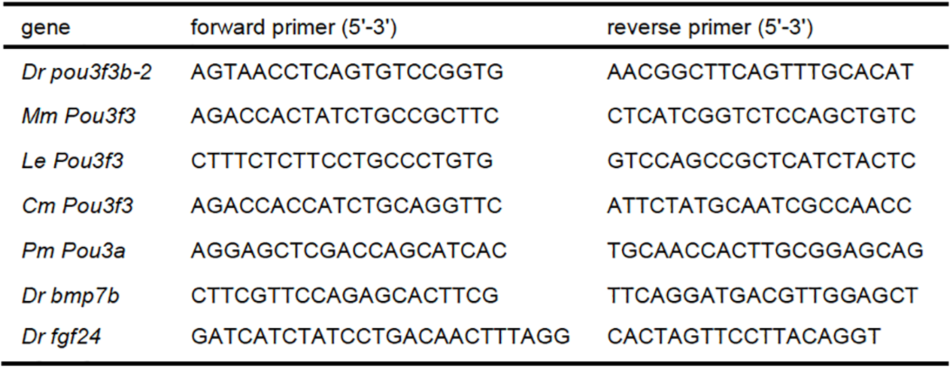
Primers for in situ probes.

**Extended Data Table 3.**
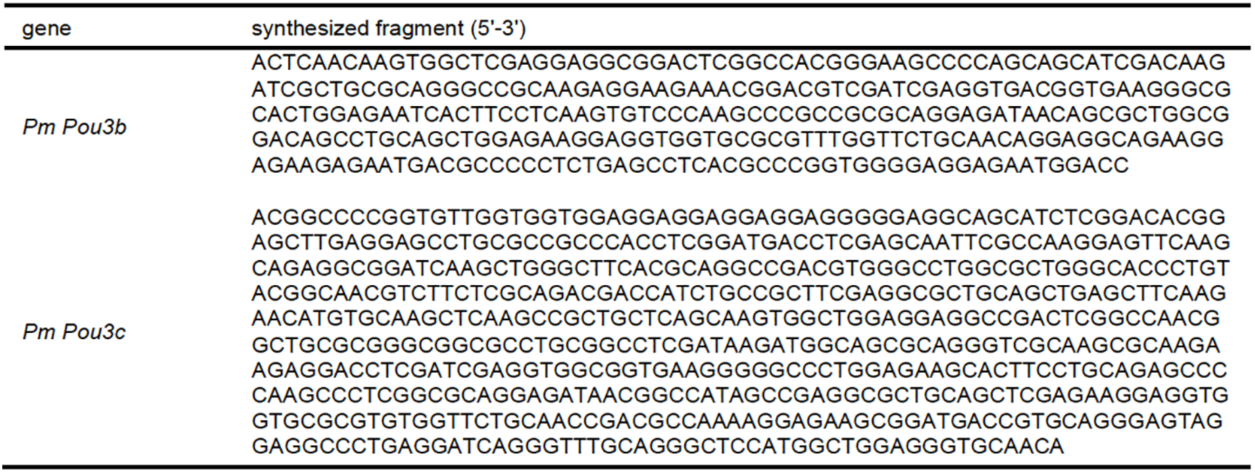
Synthesized cDNA sequences for lamprey in situ probes.

**Extended Data Table 4.**
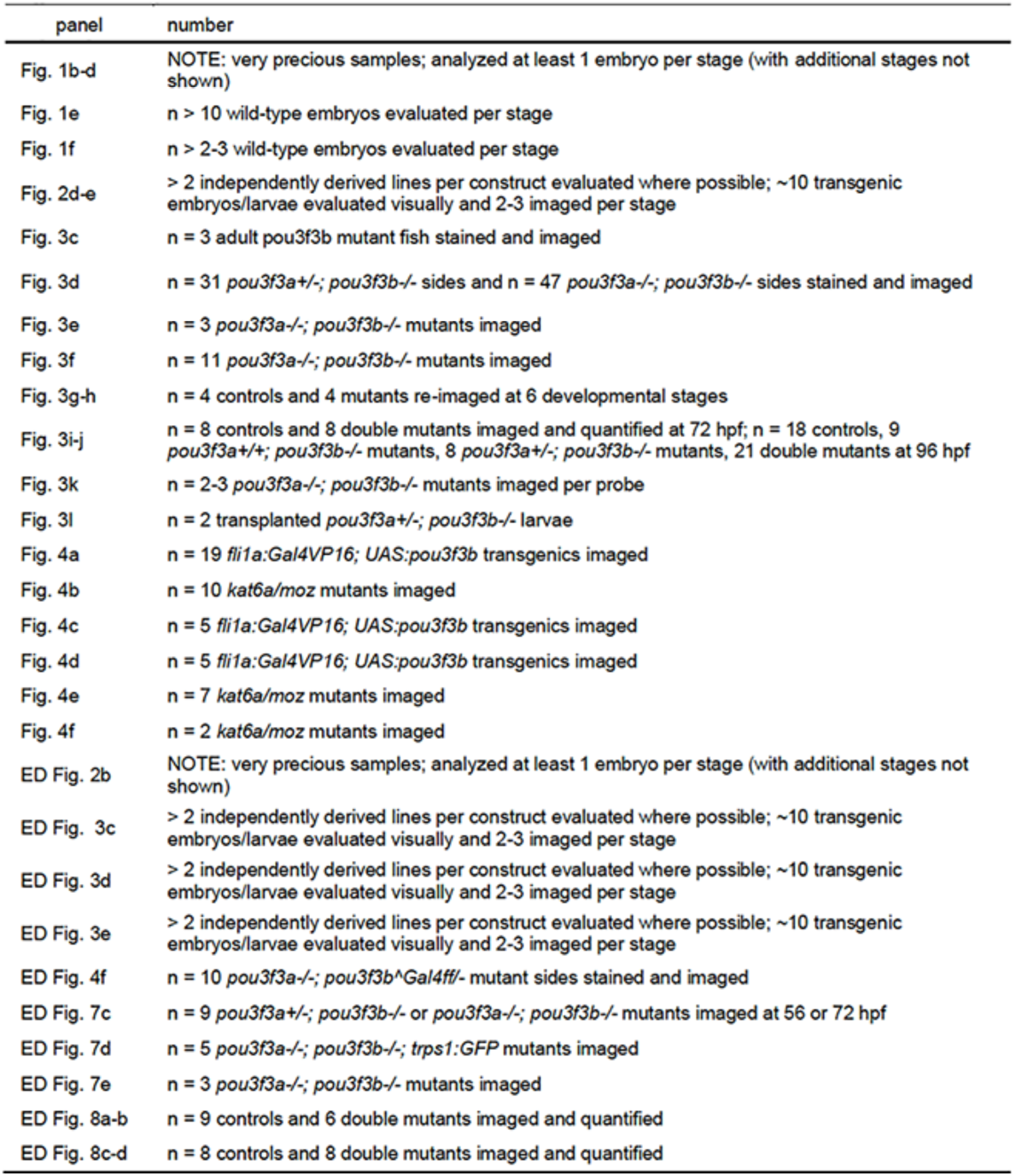
Experimental numbers.

